# The Toxin Cercosporin is a Virulence Factor for Infection of Coffee by *Cercospora coffeicola*

**DOI:** 10.1101/818328

**Authors:** A. G. C. Souza, S. Herrero, M. E. Daub

## Abstract

Brown eye spot, caused by *Cercospora coffeicola*, causes significant losses in both quality and quantity of coffee production. As many *Cercospora* spp. produce the photoactivated toxin cercosporin, this study aimed to determine the role of cercosporin in *C. coffeicola* pathogenesis by creating disruption mutants unable to produce the toxin. Six *C. coffeicola* isolates from Brazilian fields, representing organic and conventional production systems in the Minas Gerais state, were evaluated for their ability to produce cercosporin *in vitro*. Toxin production varied among isolates, ranging from 3.5 – 25.3 µM/ 5 mm mycelial plug; production was undetectable in one isolate. The *C. coffeicola* homolog of the polyketide synthase gene (*CTB1*) involved in cercosporin production was amplified using a degenerate primer strategy. The 7044 nt *ccCTB1* gene sequence was 90.3% identical to the *cnCTB1* gene in *Cercospora nicotianae* and encoded a putative protein of 2196 amino acids with 98.2% similarity and 97.5% identity to its counterpart in *C. nicotianae*. Transformation of two isolates of *C. coffeicola* with a *CTB1* disruption construct resulted in the recovery of six *ctb1* disruption mutants. All of the *ctb1* disruptants were deficient in cercosporin production. Disruption mutants did not differ significantly from the wild type for either growth or sporulation, but were significantly altered in virulence on coffee. As compared to wild type, time to lesion development was significantly increased and numbers of lesions were significantly decreased in coffee plants inoculated with *ctb1* disruption mutants. These results show that cercosporin toxin is a virulence factor for *C. coffeicola* infection of coffee.

## INTRODUCTION

Brazil is the major world producer of coffee. An important coffee disease is brown eye spot, caused by *Cercospora coffeicola*, which causes losses in both quality and quantity of production (36). Although the importance of the disease is well known, there are no cultivars resistant to the pathogen, and disease control relies primarily on fungicide sprays. The infection process of *C. coffeicola* on the coffee leaf has been studied (29), but little else is known about *C. coffeicola* pathogenesis mechanisms.

Plant pathogenic fungi use many strategies to cause disesase. One common strategy involves the production of toxins that kill host cells, allowing for tissue colonization by the pathogen (2, 25). Many *Cercospora* species produce cercosporin (23), a perylenequinone toxin. Cercosporin is light-activated, generating reactive oxygen species (ROS) such as singlet oxygen and superoxide (12, 14). The production of toxic ROS makes cercosporin almost universally toxic, with toxicity documented to host and non-host plants, bacteria, many fungi, and even mice (13). In host plants, cercosporin secreted by the fungus accumulates in membranes, leading to peroxidation of membrane lipids and death of the host cells, allowing for tissue colonization by the fungus (11, 14). The biosynthetic pathway for cercosporin production has been elucidated in *C. nicotianae* (4, 5, 6, 7, 16). The gene cluster contains genes encoding a polyketide synthase (*CTB1*; <u>C</u>ercosporin <u>T</u>oxin <u>B</u>iosynthesis 1), two *O*-methyltransferases (*CTB2, CTB3*), three oxidoreductases (*CTB5, CTB6, CTB7*), an MFS transporter (*CTB4*), and a transcriptional activator (*CTB8*). Genes involved in cercosporin production not found in the gene cluster have also been characterized in several species. These include *CFP* (<u>C</u>ercosporin <u>F</u>acilitator <u>P</u>rotein, encoding an MFS transporter) in *C. kikuchii* and *C. beticola* (3, 15) and *C. nicotianae* (31); *CRG1* (<u>C</u>ercosporin <u>R</u>esistance <u>G</u>ene, encoding a zinc cluster transcription factor) in *C. nicotianae* (8, 10); *ATR1* (encoding an ABC Transporter) in *C. nicotianae* (1, 21); and *CZK3* (encoding a MAP kinase) in *C. zeae-maydis* (26). Non-cercosporin-producing mutants of *C. nicotianae* and *C. beticola*, disrupted for genes in the biosynthetic pathway, produced fewer lesions as compared to the wild-type when these mutants were inoculated onto tobacco and sugar beet leaves, demonstrating that cercosporin is an important virulence factor in disease development on some hosts (4, 7, 16, 30, 33). By contrast, cercosporin is not produced by strains of *Cercospora arachidicola*, and there is no evidence that cercosporin plays a role in disease on peanut (18).

There are no reports on the genes involved in cercosporin production by *C. coffeicola*. It has been shown that shading reduces fungal penetration, and fewer lesions develop on shaded leaves (17). In previous studies (28), we found high variability in cercosporin production by isolates of *C. coffeicola* collected in different locations in the Minas Gerais state of Brazil. We also found a positive correlation between cercosporin production *in vitro* and virulence (28). These studies suggested that cercosporin can be an important virulence factor for *C. coffeicola* infection on coffee, and that efforts to identify genes involved in cercosporin resistance may allow for novel strategies for developing coffee with resistance to brown eye spot, by targeting toxin production or resistance.

The goal of this work was to determine the role of the cercosporin toxin in virulence of *C. coffeicola* by creating cercosporin-deficient mutants and investigating changes in disease development. Here we report the identification and sequence of a *C. coffeicola* homolog of *CTB1* (*ccCTB1*) encoding the cercosporin polyketide synthase (PKS), the recovery and characterization of cercosporin-deficient mutants by disruption of *ccCTB1*, and virulence of these mutants on coffee. Our results document an important role for cercosporin in virulence of *C. coffeicola* on coffee.

## MATERIALS AND METHODS

### Isolates

Six isolates of *C. coffeicola* differing in production of cercosporin and virulence were recovered from three Minas Gerais regions (Mata [M], Sul de Minas [S], and Triângulo [T]) and from two cropping systems (conventional [C] and organic [O]). The six isolates used were: WT-MO53, WT-SO40, WT-SC31, WT-MC56, WT-TO02, WT-TC07 (WT = wild type; M, S, T = regions; C, O = crop systems; numbers are random). The isolates were routinely cultured on “complete medium” agar or potato dextrose agar (PDA) at 25ºC in either lighted (for cercosporin production) or dark growth chambers as described (1). All analysis of variance and means comparisons were performed with SAS^®^ v. 9.1. The assay to evaluate the ability of the *ctb1* mutants to infect the coffee plants *in vivo* was conducted at the Universidade Federal de Viçosa. All other experiments were conducted at North Carolina State University.

### *Cercospora coffeicola* DNA isolation

Mycelial cultures from either wildtype or *ctb1* mutants were grown as previously described (9), and genomic DNA was isolated from lyophilized mycelium using the Fungal DNA Miniprep kit (Omega Bio-Tek, Norcross, GA, USA) following the manufacturer’s recommendations.

### Amplification of a polyketide synthase (*ccCTB1*) from *Cercospora coffeicola*

The *C. coffeicola CTB1* gene (*ccCTB1*) was amplified by PCR from an isolate showing high levels of cercosporin and virulence (WT-SO40), using first, *cnCTB1*-specific primers derived from conserved regions of the *C. nicotianae CTB1* sequence (GenBank accession number AAT69682.1), including degenerate primers derived from highly conserved polyketide synthase sequences (Ketosyntase [KS] and Acyl Transferase [AT] domains), and second, *ccCTB1*-specific primers derived from sequence information generated above. Sequence information on non-coding 5’- and 3’- end flanking regions of *ccCTB1* was also generated. PCR was performed in a 50 μl mix using Apex ^®^ Taq Polymerase (Genesee Scientific, San Diego, CA, USA) following standard procedures. The PCR products were sent to Eton Bioscience, Inc. (Research Triangle Park, NC) for sequencing, and gene sequences were assembled using the sequence analysis software Vector NTI v.10 (Invitrogen, Carlsbad, CA).

### *CTB1* gene disruption

The isolates WT-SO40 and WT-SC31, showing the highest cercosporin production and high virulence, were selected for disruption of *ccCTB1*. A split-marker disruption strategy similar to the one described previously described (1, 6, 34) was used to create *ccCTB1-*disrupted isolates. The disruption vector pCTB115 containing a hygromycin-resistance gene (*HYG*) flanked by *cnCTB1* sequences from *C. nicotianae* was used for replacement of the *ccCTB1* homolog in *C. coffeicola*. Two different overlapping PCR fragments were amplified from pCTB115 and were used as part of the split-marker strategy to disrupt *ccCTB1*. A 2.6-kb fragment containing the 5′-end sequences of *cnCTB1* and another 2.4-kb fragment containing 3′-end sequences of *cnCTB1* were amplified as previously described (6). Protoplasts were generated and transformed as previously described (19, 20, 27). The transformed cultures were selected on regeneration medium containing 125 μg/ml of hygromycin as described (1). The transformants were screened initially for lack of cercosporin production in culture according to the methodology described below. *C. coffeicola* transformants that did not produce cercosporin were verified by PCR to confirm successful disruption. Primers 3R (5’-CTCCAAGAACGTTTCGCTGT-3’) and OUTF (5’-CCATCTCATCTGCACTTCCGTTCTT-3’) were used to amplify a fragment of *ccCTB1* gene specific only to the intact WT sequence. The 3R primer was designed to hybridize inside of the *ccCTB1* region (position of the 3240 nt) putatively disrupted by the disruption-cassette, whereas the OUTF primer hybridized at the 5’ upstream region (−200 nt). The second set of primers OUTF and Hyg3 (5’GGATGCCTCCGCTCGAAGTA3’) were used to amplify a region spanning the disruption construct to confirm gene disruption the *ccCTB1* gene by the HYG gene.

### Cercosporin production assay

To assay for cercosporin production, the transformants and the wild type were grown on PDA at 25ºC under 12 h of light, conditions that induce cercosporin production in the wild type. The production was visually identified as a red pigment on the under-side of colonies 4-5 days after plating. Cercosporin production was quantified by extracting mycelial plugs in 5N KOH as described (6). The amount of cercosporin in the extract was quantified spectrophotometrically by measuring absorbance at 480, 590 and 640 nm (35). The wild type isolate used to generate the mutants as well as the other *C. coffeicola* isolates (WT-MO53, WT-MC56, WT-TO02, and WT-TC07) were used as controls. The experiment was conducted twice, each time in a randomized complete block design with three replicates (one tube = one experimental unit).

### Growth and conidia production

Growth of *C. coffeicola* wild type and *ccctb1* mutant isolates was measured on PDA with and without hygromycin (125μg/ml) by measuring the colony diameter every 3 days for 12 days. The technique of drying the mycelial mass was used to induce conidial production (29). Briefly, mycelial plugs were grown with shaking in 10 ml liquid V8 medium (20% V8) at 120 rpm, 25ºC, 12 h light. After 4 days, the cultures were transferred to open Petri dishes under fluorescent lamps, at 25ºC and with a 12-h photoperiod. After dehydration of the culture medium (approximately 4 days), 10 ml of distilled water were added to each Petri dish, the fungal colony was scratched with a glass rod, and the suspension was filtered through one layer of cheesecloth. The conidial concentration was evaluated with a haemocytometer. Each experiment was conducted twice, each time in a randomized complete design with three replicates (one plate = one experimental unit). Each experiment was analyzed separately.

### Pathogenicity of *ctb*1 disruptants on coffee

Inoculation experiments utilized two *ccctb1*-disrupted isolates generated from the WT-SC31 (MUT7 and MUT12), one transformant that was not disrupted (MUT13), and the WT-SC31. Plants of ‘Catuaí Vermelho IAC44’ (‘Catuaí’) at six months after sowing with two pairs of leaves were used for the inoculation experiments. The plants were grown in plastic bags (10 × 20 cm) containing a mixture of soil, sand, and cow manure in a proportion of 3:1:1 (v:v:v). The inoculum suspension was adjusted to 4.5 × 10^4^ conidia/mL and sprayed on both sides of four leaves per plant using a DeVilbiss sprayer. Distilled water was sprayed on the control plants. For the first 12 h after inoculation, the plants were set in a wet chamber, which was kept at 25ºC with 90 ± 5% relative humidity (RH), and continuous light (40 W grow lux lamps distributed alternately to provide light intensity of 165.3 µmol ∕ s ∕ m^2^). Subsequently, the plants were transferred to a greenhouse with natural daylight and without temperature control.

Each plant was assessed at five-day intervals from the 10^th^ to the 50^th^ day after inoculation. The evaluations measured the incubation period (IP, defined as the the number of days between inoculation and the appearance of the first symptom on a leaf), as well as lesion number and disease severity at the 35^th^ and 50^th^ day after inoculation. For each plant, the IP, lesion number and severity were calculated by averaging the values of the four inoculated leaves. The disease severity was evaluated based on a diagrammatic scale (24) with five grades: 1 (0%); 2 (>0 to 3%); 3 (>3 to 6%); 4 (>6 to 12%); and 5 (>12 to 25%) diseased area. The intermediate severity value between the lower and the upper limits of each grade was used in the statistical analysis. The contrast procedure (α=0.01) of SAS^®^ v. 9.1 was used to compare the mean of IP, number of lesions and severity at the 35^th^ and 50^th^ day after inoculation. The experiment was conducted twice, each time in a randomized complete design with seven replicates (one plant with four leaves= one experimental unit). Each experiment was analyzed separately.

## RESULTS

### Cercosporin production by wild type isolates

Cercosporin production of *C. coffeicola* wild type (WT) cultures was quantified from mycelium grown on PDA. Cercosporin production values were significantly different from each other according the Tukey test (α=0.05). Isolates WT-SO40 and WTSC31 had the highest values, 25.17 and 16.50 μM/0.5 cm mycelial plug, respectively. Isolates WT-TC07, WT-ZO53, and WT-TC02 produced 9.3, 7.5, and 4.0 μM/0.5 cm mycelial plug, respectively; production was undetectable in isolate WT-ZC56. The WT isolates with highest levels of the cercosporin production (WT-SO40 and WTSC31) were selected for disruption of the *ccCTB1* gene.

### Amplification of a cercosporin polyketide synthase gene from *C. coffeicola* (*ccCTB1*)

The full-length sequence of *ccCTB1* was determined as described in Materials and Methods by sequencing multiple overlapping PCR fragments amplified from WT-SO40 (highest cercosporin production and virulence). The size of the *ccCTB1* (GenBank accession number HQ173811.1) from *C. coffeicola* is 7044 nt, and it is 90.3% identical to *cnCTB1* from *C. nicotianae*. The size of the ccCTB1 protein in *C. coffeicola* is 2196 aa, and it is 98.2% similar and 97.5% identical to its counterpart in *C. nicotianae*. Similar to *C. nicotianae*, the *ccCTB1* gene contains a total of eight small introns. The *ccCTB1* introns ranged from 47 to 80 bp in size and are distributed throughout the gene as follows: Intron 1: 294-344; Intron 2: 649-706; Intron 3:1488-1534; Intron 4: 1687-1743; Intron 5: 1793-1839; Intron 6: 4504-4583; Intron 7: 4940-5000; and Intron 8: 6314-6368. Alignment analysis of CTB1 sequences of *C. nicotianae* (GenBank accession number AY649543) and *C. coffeicola* (GenBank accession number HQ173811.1) showed a high level of conserved amino acids between these proteins, with identity and similarity values of 97 and 93%, respectively.

### *ccCTB1* gene disruption and evaluation of cercosporin production

The *C. coffeicola ccCTB1* gene showed high homology to *cnCTB1* in *C. nicotianae.* Thus the vector pCTB115 used to disrupt *CTB1* from *C. nicotianae* (6) was used for disruption of the *ccCTB1* gene in *C. coffeicola* using a split marker approach. The two wild type isolates with the highest levels of cercosporin production, WT-SO40 and WT-SC31, were selected for *ccCTB1* disruption. Transformants were selected for hygromycin (HYG) resistance. HYG-resistant isolates were then screened for lack of visible red pigmentation (cercosporin) when grown on PDA. To ensure that transformants did not contain a mix of disrupted and wild type nuclei, isolates were transfered six times on PDA amended with 125 μg/ml of hygromycin before quantifying cercosporin production and confirming disruption by PCR.

Cercosporin production was assayed by extraction in KOH and quantification by absorption spectroscopy. Cercosporin is readily detectable by a green-color when extracted in 5N KOH (Fig. 1), and absorption spectroscopy showed the presence of typical peaks at 480, 590 and 640 λ that were not observed in isolates that failed to produce cercosporin (data not shown). Out of 126 HYG-resistant transformants obtained for both isolates, nine (4/111 transformants of WT-SO40; 5/15 transformants of WT-SC31) did not have detectable levels of cercosporin as assayed by KOH extraction (Fig. 1). Cercosporin production by one randomly selected cercosporin-producing transformant from each parent (MUT5 from WT-SO40 and MUT13 from WT-SC31) was quantified and shown to be similar to their wild type parents; WT-SO40 and MUT 5 produced 43.5 and 40.5 μM/0.5 cm mycelial plug, respectively, and WT-SC31 and MUT13 produced 26.5 and 24.3 μM/0.5 cm mycelial plug, respectively.

**Figure 1.**
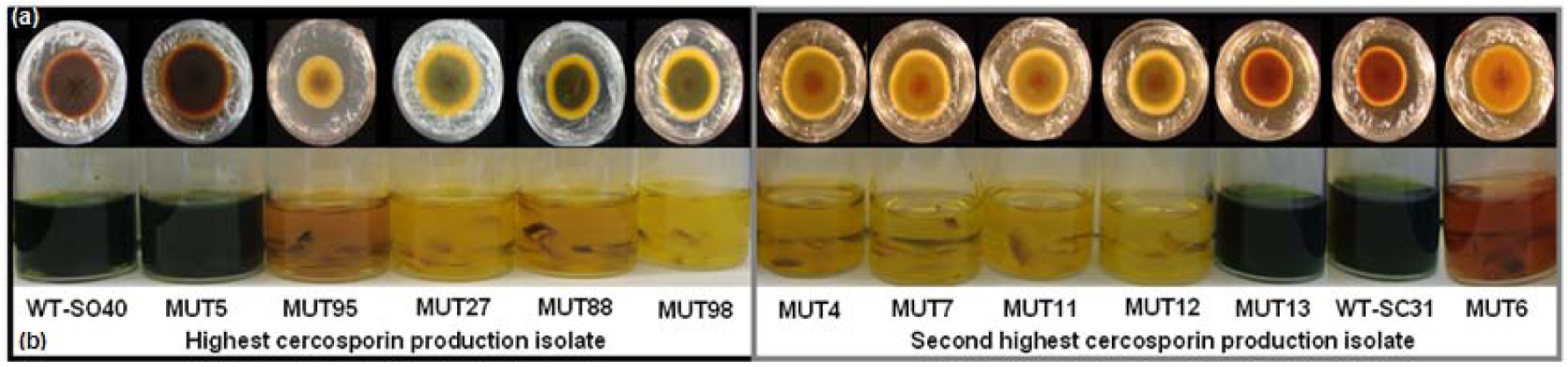
Cercosporin production by wild type (WT) *C. coffeicola* isolates and transformants (MUT) following transformation with the *ctb1* disruption construct. Transformants were recovered from transformation of the two highest cercosporin-producing wild type isolates: WT-SO40 (MUT5, MUT95, MUT27, MUT88, and MUT98) and WT-SC31 (MUT7, MUT11, MUT12, MUT13, and MUT6). **(a)** Colonies grown in PDA with HYG (12 days, 25ºC, 12h light). Red pigmentation indicates certosporin production. **(b)** Extraction of cercosporin from mycelial plugs using 5N KOH. Dark green extract indicates presence of cercosporin. WT-SO40, WT-SC31 and two transformants, MUT5 and MUT13, produced cercosporin whereas all remaining transformants (MUT95, MUT27, MUT88, MUT98, MUT4, MUT7, MUT11, MUT12, MUT6) did not produce cercosporin.

### Confirmation of *ccctb1* disruption by PCR

The nine non-cercosporin-producing transformants and two selected cercosporin-producing transformants were analyzed by PCR to confirm targeted disruption of *ccCBT1* gene of *C. coffeicola*. Primers 3R (in the *ccCTB1* ORF) and OUTF (upstream of coding sequence) were used to amplify a fragment of *ccCTB1* gene specific only to the intact WT sequence. The second set of primers (OUTF and Hyg3) was used to amplify a region spanning the disruption construct to a genomic region outside the construct to confirm homologous integration and gene disruption the *ccCTB1* gene by the HYG gene. Of the wild type parent strains, nine transformants lacking cercosporin production, and two cercosporin-producing transformants analyzed, only the two wild type strains showed the presence of the intact *ccCTB1* gene (Fig. 2). Six of the nine cercosporin-deficient transformants (MUT4, MUT7, MUT12, MUT88, MUT95 and MUT98) were confirmed to lack the intact *ccCTB1* gene and also contain the expected band confirming *ccCTB1* disruption. In addition, three cercosporin-deficient transformants (MUT6, MUT11 and MUT27) and two cercosporin-producing transformants (MUT5, MUT13) lacked a detectable wild type copy; however, the disruption of *ccCTB1* could not be confirmed for these isolates. These results may indicate rearrangements during the integration process.

**Figure 2.**
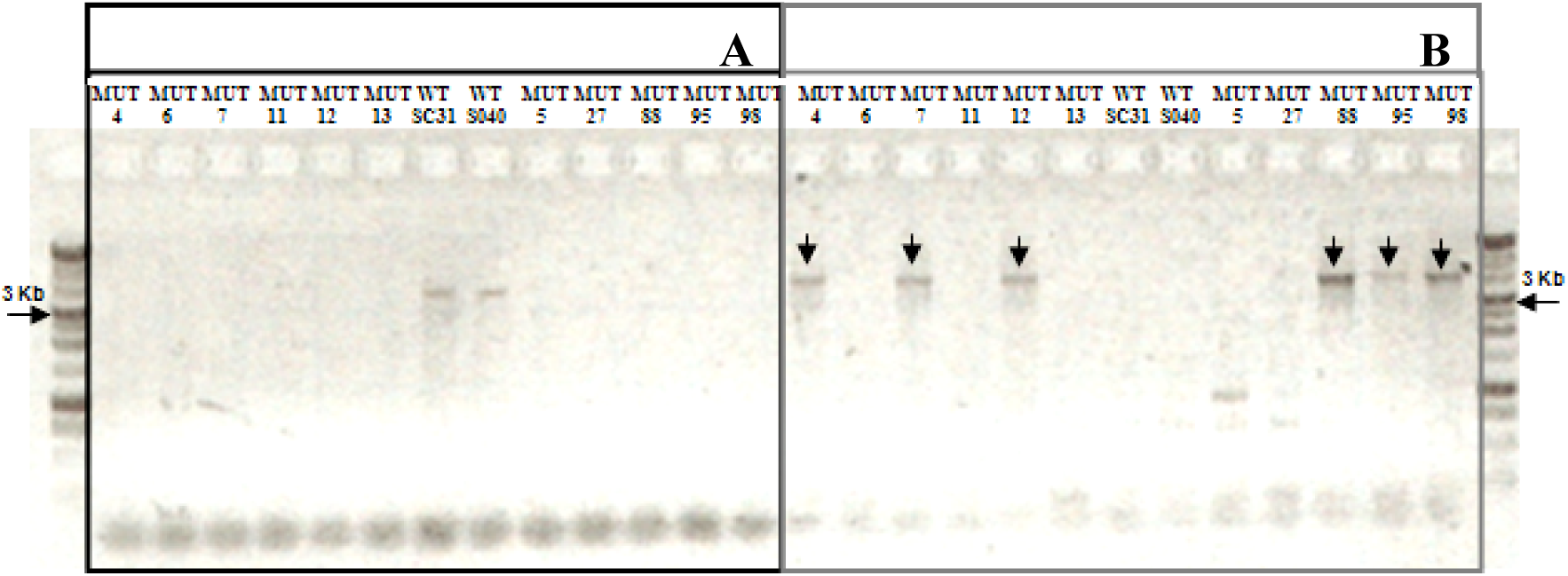
Molecular analysis of *C. coffeicola* wild type (WT) and transformants transformed with the *ctb*1 disruption construct (MUT). Protoplasts were transformed as described in Materials and Methods. Regenerated colonies were selected for HYG resistance, and screened for lack of cercosporin production. Two sets of primers, OUTF and 3R [A] or OUTF and Hyg3 [B], were used to identify, respectively, the WT (3.5Kb) or disrupted (4.0Kb) copies of the *ccCTB1* gene in isolates. Panel A shows amplification of the WT gene in SC31 and SO40. Amplified bands in panel B indicate presence of the disrupted gene. Arrows indicate disrupted mutants. MUT5, MUT27, MUT88, MUT95 and MUT98 isolates came from WT-SO40; MUT4, MUT6, MUT7, MUT11, MUT12 and MUT13 isolates came from WT-SC31.

### Growth and conidia production of *ccctb1* disrupted isolates

Wild type, the six confirmed *ccctb1*-disrupted transformants, and selected non-disrupted transformants were analyzed for growth rate in culture and conidial production. As expected, neither of the wild type isolates (WT-SO40 and WT-SC31) grew on PDA supplemented with hygromycin. There were no significant differences in growth on PDA medium between the wild type isolates and any of the transformants, either the *ccctb1* disruptants or the non-disrupted transformants (Table 1).

**TABLE 1.** Average radial growth (diameter in cm) after 12 days of wild type parent isolates (WT), *ccctb1-*disrupted transformants (**), and non-disrupted transformants (^x^) grown on potato dextrose agar (PDA) and PDA supplemented with hygromycin (PDA+HYG), in two different experiments

| Wild Type<br>Parent | Isolate | First experiment |  | Second experiment |  |
| --- | --- | --- | --- | --- | --- |
|  |  | PDA* | PDA+HYG* | PDA* | PDA+HYG* |
| WT-SO40 | WT-SO40 | 1.11 A | 0.00 B | 1.80 A | 0.00 B |
|  | MUT5<sup>x</sup> | 1.14 A | 1.23 A | 1.91 A | 2.00 A |
|  | MUT95** | 1.26 A | 0.95 A | 1.92 A | 1.30 A |
|  | MUT88** | 1.47 A | 1.18 A | 2.00 A | 1.80 A |
|  | MUT98** | 1.59 A | 1.43 A | 2.35 A | 2.03 A |
|  | MUT27<sup>x</sup> | 1.64 A | 1.31 A | 2.22 A | 2.11 A |
| WT-SC31 | WT-SC31 | 1.28 A | 0.00 B | 1.79 A | 0.00 B |
|  | MUT13<sup>x</sup> | 1.35 A | 1.22 A | 1.84 A | 1.98A |
|  | MUT6<sup>x</sup> | 1.44 A | 1.36 A | 2.19 A | 2.13 A |
|  | MUT7** | 1.52 A | 1.40 A | 2.09 A | 2.09 A |
|  | MUT4** | 1.53 A | 1.44 A | 2.10 A | 2.22 A |
|  | MUT11<sup>x</sup> | 1.55 A | 1.54 A | 2.28 A | 2.23 A |
|  | MUT12** | 1.55 A | 1.51 A | 2.20 A | 2.22 A |

\*Values with different letters in the same column of a wild type isolate indicate significantly different growth (Tukey's test, $\alpha = 0.05$ ).

For sporulation, differences were found in numbers of spores produced between isolates in both experiments (P < 0.0001). However, there was no consistent difference between the wild type parents and the *ccctb1* disruptants across both experiments (Table 2).

**TABLE 2.** Average sporulation (conidia/ml) of wild type (WT), *ctb1*-disrupted-transformants (**), and non-disrupted transformants (^x^) in two experiments

| Wild Type |  |  |  |
| --- | --- | --- | --- |
| Parent | Isolate | First experiment* | Second experiment* |
| WT-SO40 | WT-SO40 | 262500 AB | 145000 A |
|  | MUT5<sup>x</sup> | 232500AB | 185000A |
|  | MUT27<sup>x</sup> | 215000 B | 215000 A |
|  | MUT95** | 27500 C | 145000 A |
|  | MUT88** | 18000 C | 125000 A |
|  | MUT98** | 13000 C | 175000 A |
| WT-SC31 | MUT12** | 292500 A | 170000 C |
|  | MUT7** | 282500 AB | 217500 C |
|  | MUT6<sup>x</sup> | 225000 ABC | 342500 B |
|  | MUT13<sup>x</sup> | 185000C | 292500BC |
|  | MUT11<sup>x</sup> | 165000 C | 485000 A |
|  | WT-SC31 | 16750 D | 362500 B |
|  | MUT4** | 14250 D | 402500 AB |

\*Values with different letters in the same column of a wild type isolate indicate significantly different sporulation (Tukey's test, $\alpha = 0.05$ ). For the statistical analysis, the sporulation values were transformed to log (sporulation).

### Pathogenicity of *ccctb1* disruptants on coffee

To determine the importance of cercosporin in *C. coffeicola* pathogeneis on coffee, WT-SC31, two independent *ccctb1* disrupted mutants from WT-SC31 (MUT7, MUT12), and a non-disrupted, cercosporin-producing transformant from WT-SC31 (MUT13) were inoculated onto coffee. Incubation period (IP), lesion numbers, and disease severity ratings were determined. In both experiments, medium values of the IP and SEV35 and SEV50 (disease severity at the 35^th^ and 50^th^ day after inoculation) were not stastically different between the cercosporin-producing wild type WT-SC31 and the non-disrupted cercosporin-producing MUT13 transformant (Table 3), showing that the transformation procedure did not affect virulence. Differences were seen between the two cercosporin-producing strains (WT-SC31 and the non-disrupted MUT13) and the disrupted strains (MUT7, MUT12) that do not produce cercosporin. The incubation periods (IP) associated with the cercosporin-producer isolates (WT-SC31 and MUT13) were statistically different (P < 0.01) from the non-cercosporin producers (MUT7 and MUT12) in both experiments. The average values of the IP of the two cercosporin-producing isolates (WT-SC31 and MUT13) were 30.3 and 32.3 days whereas IP for the two disrupted non-producing mutants (MUT7 and MUT12) were 38.3 and 41.3 days on the first and second experiments, respectively (Table 3). The averages of the number of lesions (NL) produced on leaves were also significantly different between cercosporin-producing and non-producing isolates. NL associated with the cercosporin-producing isolates was significantly higher (P<0.01) than for the non-producers at the 35^th^ and 50^th^ days after inoculation in both experiments (Table 3). Variation in severity values was high, limiting statistical significance.

**TABLE 3.** Number of lesions (NL), severity (SEV), and incubation period (IP) of brown eye spot on coffee leaves caused by *Cercospora coffeicola* isolates. Leaves were inoculated with wild type (WT-SC31), a cercosporin-producing non-disrupted transformant (MUT13), and two *ccctb1* disupted mutants (MUT7, MUT12) unable to produce cercosporin. NL and SEV were evaluated at 35 (NL35, SEV35) and 50 (NL50, SEV50) days after inoculation. Each value is the mean of three replications per experiment in two independent experiments (E1 and E2).

| Isolate | Cerc<sup>1</sup> | NL35 |  | NL50 |  | SEV35 |  | SEV50 |  | IP |  |
| --- | --- | --- | --- | --- | --- | --- | --- | --- | --- | --- | --- |
|  |  | E1 | E2 | E1 | E2 | E1 | E2 | E1 | E2 | E1 | E2 |
| **WT-SC31** | + | 4.29 | 1.78 | 10.29 | 3.03 | 0.95 | 0.86 | 1.45 | 1.12 | 31.43 | 30.30 |
| **MUT13** | + | 6.00 | 0.63 | 7.29 | 0.82 | 1.38 | 0.53 | 1.39 | 0.70 | 29.17 | 34.29 |
| **MUT7** | - | 1.61 | 0.21 | 3.95 | 0.37 | 0.43 | 0.63 | 1.18 | 0.75 | 39.29 | 43.18 |
| **MUT12** | - | 1.32 | 0.56 | 3.79 | 0.82 | 0.79 | 0.25 | 1.02 | 0.37 | 37.32 | 39.34 |
| **Cerc+ × Cer-\*** |  | 0.0012 | 0.0002 | 0.0030 | 0.0003 | 0.0148 | 0.062 | 0.13 | 0.0188 | 0.0083 | 0.0002 |

<sup>1</sup> Cercosporin production
\*Significance (P-values) of a statistical contrast that tested whether cercosporin producer isolates (Cerc+) were different from non-producer isolates (Cerc-) regarding each variable in each experiment.

## DISCUSSION

Many species of plant pathogenic fungi produce toxins that act as virulence factors and help parasitize their hosts (2, 25). Many *Cercospora spp.* produce the light-activated photosensitizing toxin cercosporin (14, 23). Mutants of *C. nicotianae*, *C. kikuchii*, and *C. zeae-maydis* have been generated to confirm the role of cercosporin as a virulence factor (3, 6, 26). Our epidemiological studies suggested that cercosporin is involved in virulence of *C. coffeicola* on coffee, as we found a positive correlation between cercosporin production and virulence (28). To conclusively show the relatedness between cercosporin production and virulence of *C. coffeicola*, we isolated a *C. coffeicola* polyketide synthase gene homologous to the <u>C</u>ercosporin <u>T</u>oxin <u>B</u>iosynthesis gene 1 (*CTB1*) in *C. nicotianae*. We then determined *ccCTB1*’s role in fungal cercosporin biosynthesis by generating isolates lacking this gene.

In our study, sequence analysis of ccCTB1 in *C. coffeicola* revealed strong similarity to the cnCTB1 protein involved in cercosporin production by *C. nicotianae*. The ccCTB1 sequence was also similar to other polyketide synthase (PKS) sequences of other fungi, including the ALB1 protein involved in conidial pigment biosynthesis in *Penicillium marneffei* and *Aspergillus fumigates* with 65% (accession number XM002147681) and 67% (accession number XM751002) identity, respectively. As with the *C. nicotianae* CTB1, the *C. coffeicola* CTB1 contains five catalytic domains: a keto synthase (KS), an acyltransferase (AT), a thioesterase (TE), and two acyl carrier protein (ACP) domains, involved in the initial steps of cercosporin synthesis (6). Other genes, including CTB3 (encoding a dual methyltransferase/monooxygenase), CTB4 (encoding a putative membrane transporter), CZK3 (gene that regulates cercosporin biosynthesis, fungal development, and pathogenesis), and CFP (encoding a membrane transport protein required for secretion of cercosporin), are also involved with cercosporin production, confirming that cercosporin biosynthesis is a process that involves multiple genes and signals (3, 6, 7, 16, 26).

We conducted gene disruption experiments and were successful in isolating disrupted transformants for the *ccCTB1* gene that do not produce cercosporin. Analysis of cercosporin production by transformants showed a direct correlation between cercosporin production and disruption of *ccCTB1*; transformants confirmed by PCR as disruption mutants did not produce cercosporin. These results confirm that the *ccCTB1* gene is required for cercosporin biosynthesis in *C. coffeicola*. These results provide the first report of a gene involved in the cercosporin biosynthesis pathway in *C. coffeicola*.

The cercosporin-non producing *ccctb1* disruption mutants were evaluated for other traits important in disease development including growth and sporulation. In our study, disruption of *ccCTB1* did not affect growth *in vitro.* A similar result was reported when *ctb4*- and *ctb3*-disrupted isolates of *C. nicotianae* were compared; disrupted isolates produced less cercosporin although they had a similar growth as the wild type (7, 16). Similarly, disruption of the CFP gene of *C. kikuchii* led a loss of 95% of the cercosporin production as compared to the wild type isolate with no change in growth (32). *C. kikuchii* UV-generated mutants unable to produce cercosporin grew more than the wild type on solid medium, although the growth on liquid medium did not differ (32). In *C. zeae-maydis*, disruption isolates for the CZK3 gene that are defective in cercosporin production had higher vegetative growth than the wild type (26). Overall growth differences do not appear to correlate with production or lack of production of cercosporin, and may be due to the specific genes defective in these mutants.

*Cercospora spp.* sporulate profusely on the lesions to generate secondary inoculum that accounts for biological fitness and survival. There are reports of non-cercosporin producing *C. kikuchii* and *C. zeae-maydis* isolates having reduced sporulation (3, 26, 32). However, in other studies *C. nicotianae ctb1* and *ctb3* mutants unable to produce cercosporin sporulated similarly to the wild type isolate (6, 16). We did not find a significant difference in sporulation between cercosporin-producing and non-producing isolates of *C. coffeicola*. Considering the relevance of sporulation on disease progress on the field, it would be interesting to quantify both cercosporin production and sporulation on the host tissue to verify if there is any relation between them.

The role of the cercosporin production in the ability of *Cercospora* spp. to parasitize plants has been studied for different pathosystems (3, 4, 6, 16, 26). In our study, we found that the cercosporin-producing isolates produced more lesions as compared to the non-producing disruptants when these isolates were inoculated onto coffee leaves. In addition, incubation period was also significantly longer for the non-cercosporin producing disruptants. As secondary cycles are important in the epidemics of policyclic diseases, such as brown eye spot, changes in incubation period would alter the course of epidemics. Our results are similar to studies in other systems where mutants of *C. nicotianae* unable to synthesize cercosporin were shown to produce fewer lesions as compared to the wild type when these mutants were inoculated onto tobacco leaves (4, 16). In our study, numbers of lesions could distinguish the virulence of cercosporin-producing from the non-producing isolates. However, disease severity measurements did not always correlate with cercosporin production. Severity was visually estimated, thus errors and variability can be larger than lesion numbers and obscure any statistical difference. The use of diagrammatic scales is an effective method to evaluate the disease severity, but the occurrence of visual errors is frequently observed with the use of this technique (22). It is also possible that cercosporin can be more important for *C. coffeicola* infection (thus, the difference in the number of lesions) than for the colonization of leaves (thus, the partial differences in the severity values). Our results confirm our epidemiological and correlative data on the importance of cercosporin in *C. coffeicola* pahtogenicity (28). Our study also demonstrates there are additional mechanisms and genes that are involved in *C. coffeicola* pathogenicity because mutants that were unable to produce cercosporin were able to parasitize the coffee leaf and produce lesions.

In summary, we isolated and characterized a polyketide synthase gene from *C. coffeicola* responsible for production of cercosporin. Using gene-disruption technology, we created disrupted mutants unable to produce cercosporin. Analysis of disease development on coffee showed that the mutants produced fewer lesions with a longer incubation period. Both of these factors will strongly influence the progress and spread of the disease in the field, underscoring the importance of cercosporin in brown eye spot. This is the first report where molecular genetic manipulation of *C. coffeicola* has been used to define a critical pathogenicity factor. In addition to confirming the long-held hypothesis of the importance of cercosporin in brown eye spot, the results from this study provide fundamental information needed to address the development of novel disease control strategies. Possible control strategies include the development of fungicides that block the cercosporin synthesis, engineering coffee to silence cercosporin synthesis, or engineering or breeding coffee for toxin resistance. Knowledge gained from this work advances our understanding of *C. coffeicola* and its interactions with coffee plants, and may provide practical uses for our knowledge to better control brown eye spot.

## ACKNOWLEDGMENTS

We thank Dr. Luis Maffia, Departamento de Fitopatologia, Universidade Federal de Viçosa, Viçosa, MG 36570000, Brazil, for support and assistance with the inoculation experiments. Support for this research was provided by CNPq/NCSU/UFV and FAPEMIG.

